# NeuroVault.org: A web-based repository for collecting and sharing unthresholded statistical maps of the human brain

**DOI:** 10.1101/010348

**Authors:** Krzysztof J. Gorgolewski, Gael Varoquaux, Gabriel Rivera, Yannick Schwarz, Satrajit S. Ghosh, Camille Maumet, Vanessa Sochat, Thomas E. Nichols, Russell A. Poldrack, Jean-Baptiste Poline, Tal Yarkoni, Daniel S. Margulies

## Abstract

Here we present NeuroVault — a web based repository that allows researchers to store, share, visualize, and decode statistical maps of the human brain. NeuroVault is easy to use and employs modern web technologies to provide informative visualization of data without the need to install additional software. In addition, it leverages the power of the Neurosynth database to provide cognitive decoding of deposited maps. The data are exposed through a public REST API enabling other services and tools to take advantage of it. NeuroVault is a new resource for researchers interested in conducting meta and coactivation analyses.

## Introduction

Noninvasive neuroimaging techniques such as MRI and PET have enabled unprecedented insight into the localization of various functions in the human brain. As the number of studies using such techniques continues to grow exponentially, the challenge of assessing, summarizing, and condensing their findings poses ever greater difficulty. Even though a single study can take years to conduct, cost hundreds of thousands of dollars, and require the effort of dozens of highly trained scientists and volunteers, the output is usually reduced to an academic article, and the original data are rarely shared (Poline et al., 2012). Unfortunately due to the historical legacy of reporting knowledge in a written form (of an academic paper), the final documented results consist mostly of subjective interpretation of data with very little machine readable information. While the introduction of common stereotaxic spaces (e.g., Talairach and MNI305) has provided an initial framework for a standard of reporting activation locations, to subsequently enable meta analyses, there are several issues with this coordinate based strategies. First, peak coordinates are not able to fully describe the 3D shape and extent of a suprathreshold volume on a statistical map. Many papers use figures (2D or 3D) to present these statistical maps, but authors must decide which aspects of the 3D data cube to show. To fully explore all layers of the data one would need to be able to interrogate it in an interactive fashion. Furthermore, published figures are not machine readable, and researchers that are interested in comparing their own results with published literature are forced to manually reconstruct regions of interest (ROIs) using spheres placed at the limited reported activation locations.

A second issue is the difficulty of putting one’s results in the context of other studies. The overwhelming number of brain imaging results published each year makes manual comparison both unfeasible and prone to bias. There are attempts to automatically aggregate knowledge across large sets of neuroimaging studies. For example, Neurosynth (Yarkoni et al., 2011) is a meta analysis database that collects coordinates of activation foci from published papers and generates topic maps based on the spatial distribution of those coordinates. Such maps can aid in interpretation of new results. However, comparing a new result to a set of topic maps has so far not been implemented in a user friendly fashion.

Finally, and most importantly, making meta analytic inferences using only peak coordinates (or statistically thresholded maps) is problematic. It is easy to imagine a subthreshold effect that is consistent across many studies. Such an effect would not be picked up by existing meta analysis methods (Laird et al., 2005; Yarkoni et al., 2011) because it would never be reported in the tables of peak coordinates. Considering how underpowered most human neuroscience studies are, this situation is not that unlikely. Discarding information that is below threshold in this fashion is akin to not publishing null results (Rosenthal, 1979), a dangerous practice that creates a publication bias skewing our perception of accumulated knowledge.

Using fully unthresholded statistical maps instead of solely peak coordinates would provide a significant advance in meta analytic power. Coordinate based meta analysis (CBMA) methods show only modest overlap with image based meta analysis (IBMA; meta analysis based on unthresholded statistical maps) methods and are less powerful (Salimi Khorshidi et al., 2009). However, IBMA methods struggle with access to the data. Peak coordinates are easier to obtain and share because coordinate tables are an integral component of traditional neuroimaging papers, whereas very few papers provide links to unthresholded statistical maps (usually by an ad hoc means such as the author’s web site).

NeuroVault.org is an attempt to solve these problems. It is a web based repository that makes it easy to deposit and share statistical maps. It provides attractive visualization and cognitive decoding of the maps that can improve collaboration efforts and readability of the results. At the same time, it also provides an API for methods researchers to download the data, perform powerful analyses, or build new tools.

## Results

In the following section we describe the architecture and features of NeuroVault and present two example analyses.

### Platform

One of the key features of NeuroVault is the ease of uploading and sharing statistical brain maps. Figure 1 presents a schematic overview of the platform. After logging in, users can upload a broad range of neuroimaging images and associated metadata. These data are then immediately accessible (subject to user controlled privacy settings) via both an interactive HTML based interface, and a comprehensive RESTful web API that facilitates programmatic interoperability with other resources. In the following sections, we discuss different aspects of the platform.

**Figure 1.**
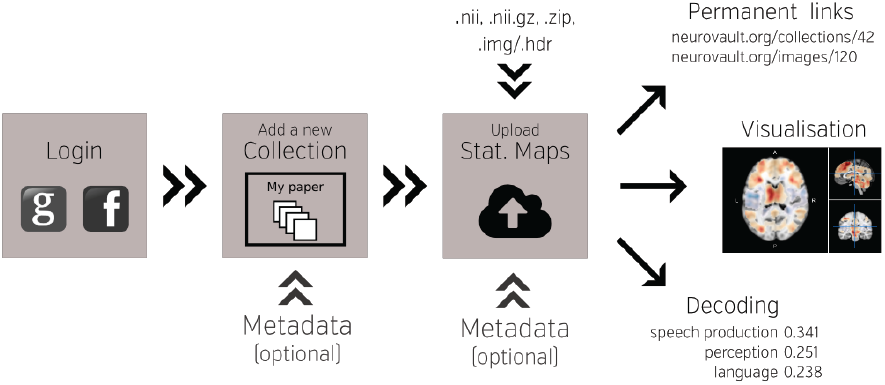
Schematic overview of the NeuroVault platform. To begin working with NeuroVault, users are asked to create an account or log in using their Facebook or Google account. After login, the user creates a collection (representing a paper or a study). At this stage, users can provide a DOI pointing to a paper associated with the collection and/or fill in a number of fields describing the study (see Supplementary Table 1 for details). This additional information is, however, optional. After the collection is created, users can add images. This can be done one by one or in bulk by uploading whole folders. Again, there is an option to add more metadata describing the images. The process of creating a collection and uploading statistical maps to NeuroVault takes only 5-10 minutes. When the maps are uploaded, users can start benefiting from permanent link to their results, interactive web based visualization, and real time image decoding.

#### Image upload

The NeuroVault upload process emphasizes speed and ease of use. Users can rely on existing social media accounts (Google or Facebook) to log in, and can upload individual images or entire folders (see Figure 1). Users can arrange their maps into collections or to group them with tags. Each collection and statistical image in NeuroVault gets a permanent link (URL) that can be shared with other researchers or included in papers or other forms of publication (blogs, tweets etc.). Users can specify whether each collection is public or private. The latter have a unique obfuscated URL that is not discoverable on the NeuroVault website, and thus are accessible only by whomever the owner decides to share the URL with. The option of creating private collections gives users freedom to decide who can access their data, and can facilitate a scenario in which a collection is shared privately during the pre publication peer review process and then made public upon acceptance of a manuscript. Using a third party (such as NeuroVault) to share data that are part of the peer review process eliminates concerns about the reviewers’ anonymity. Even though we opted to minimize the required amount of metadata^1^ for collections and statistical maps (to streamline the process) we give users an option to provide more information to maximize the usability of maps (see Supplementary Tables 1 and 2). Most importantly, we provide ability to link a collection to a paper via a DOI to promote the associated paper and facilitate meta analysis.

#### Data types

NeuroVault is able to handle a plethora of different types of brain maps as long as they are represented as 3D NIFTI files in MNI space. This includes Z or T maps derived from task based, resting state fMRI and PET experiments as well as statistics derived from analyses of structural data (e.g., Voxel Based Morphometry; VBM). In addition, results from electroencephalography (EEG) and magnetoencephalography (MEG) experiments can be used with NeuroVault as long as they are converted to NIFTI volumes through source localization (Phillips et al., 2002). NeuroVault can also handle mask files (for describing ROIs), label maps (a result of parcellation studies), posterior probability maps (coming from Bayesian methods; (Woolrich et al., 2004)), weight maps (coming from multivariate pattern analysis methods; (Haxby, 2012)), and group level lesion maps (from clinical studies). In addition, NeuroVault is able to automatically extract some metadata from SPM.mat files and FEAT folders if they are uploaded along with the statistical maps. NeuroVault also supports FSL brain atlas file format (NIFTI file with a side car XML file). When users upload such data the parcel labels are exposed through the user interface and the API (the API provides the ability to query atlases by a set of coordinates or a region name).

#### User interface

NeuroVault is designed to provide intuitive, interactive visualization of uploaded images. Each image is assigned its own unique URL with an embedded JavaScript 2D/3D viewer. In contrast to traditional, static figures in published articles, users can dynamically interact with images — adjusting statistical thresholds, selecting different color maps, and loading additional brain volumes into the viewer for comparison. Using two embedded open source JavaScript viewers (Papaya https://github.com/rii-mango/Papaya and pycortex https://github.com/gallantlab/pycortex), users can interrogate the data both in the volumetric space as well as on the surface (see Figure 2). Both viewers work inside modern web browsers and do not require any additional software to be installed. In addition to the visual representation of the volume, each page also displays any metadata associated with that image (e.g., experimental contrast, statistic type, etc).

**Figure 2.**
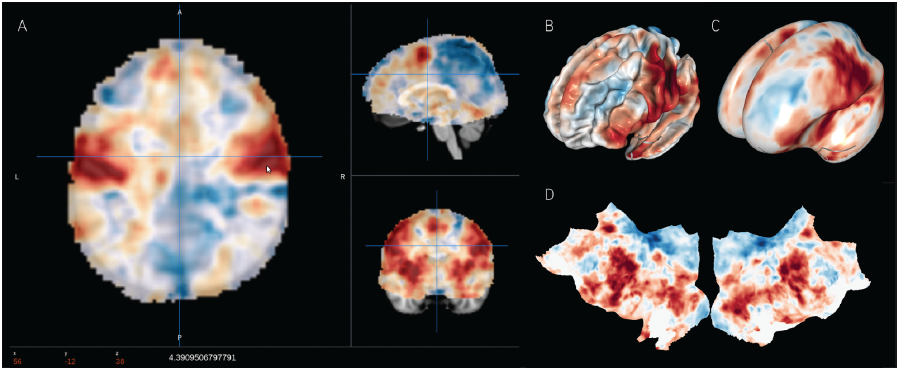
Visualization options available in NeuroVault. The user can choose to interactively interrogate the images using 2D volumetric view (A), 3D fiducial view (B), 3D inflated view (C) or a flattened cortical surface map (D).

#### Interoperability

A major goal of NeuroVault is to directly interoperate with other existing web based neuroimaging resources, ensuring that users can take advantage of a broad range of computational tools and resources without additional effort. There are two components to this. First, in cases where other relevant resources implemented a public API, NeuroVault can provide a direct interface to those resources. For example, at the push of a single button, each map deposited in NeuroVault can be near instantly “decoded” using Neurosynth (see Figure 3). In the time of 1 - 2 seconds, the uploaded image is analyzed for its spatial correlation with a subset of the concept based meta analysis maps in the Neurosynth database. The user is then presented with a ranked, interactive list of maximally similar concepts, providing a quantitative, interactive way of interpreting individual statistical images that is informed by a broader literature of nearly 10,000 studies. Second, NeuroVault exposes its own public RESTful web API that provides fully open programmatic access to all public image collections and enables direct retrieval and filtering of images and associated metadata (see http://neurovault.org/api docs for detailed description). This feature allows other researchers to leverage NeuroVault data in a broad range of desktop and web applications. To maximize the impact of data stored in NeuroVault the access to the API is unrestricted, does not require any terms of use agreements, and the data itself is distributed under the CC0 license (http://creativecommons.org/publicdomain/zero/1.0).

**Figure 3.**
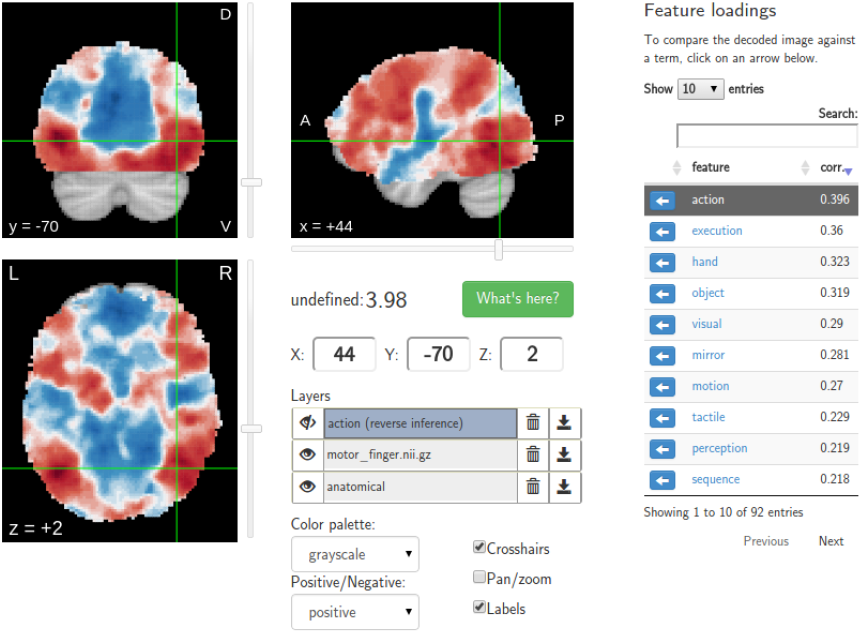
Results of the Neurosynth decoding of a statistical map obtained through NeuroVault API. Users are able to interactively compare their maps with Neurosynth topic maps.

#### Accessibility

Another advantage of depositing statistical maps in NeuroVault is the increase in longevity and impact of one’s research outputs. By providing a free, publicly accessible, centralized repository of whole brain images, NeuroVault has the potential to increase the flow of data between different researchers and lab groups. Maps deposited in NeuroVault can be used by other researchers to create detailed region of interests for hypothesis driven studies or to compare results of replications. However, one of the most interesting cases of reusing statistical maps from previous studies is image based meta analysis. Researchers wanting to perform meta analyses can obtain the statistical maps from NeuroVault and perform annotation using various external tools/platforms such as BrainMap(manual annotation; (Laird et al., 2005)), BrainSpell^2^ (crowd sourced annotation; http://brainspell.org) or NeuroSynth (automatic annotation; (Yarkoni et al., 2011)). It is worth noting that so far meta analysis in neuroimaging have rarely been performed based on labels and annotation provided by the study authors, and thus we feel outsourcing data annotation is the best current approach. Here we present a proof of concept meta analysis based on NeuroVault data collected to date. It gives a taste of the potential this platform provides for aggregating knowledge about the human brain.

### Meta-analysis using the NeuroVault data

At the time of submitting this publication, there were 135 non empty public collections (53 of them associated to a publication; for up to date stats see http://neurovault.org/collections/stats) comprised of 692 images labeled as Z, T, or F statistics. Out of these, we removed 14 outliers, and selected 678 maps to perform proof of concept analyses. The outliers were detected by using a PCA on all the statistical maps (Fritsch et al., 2012). We found wrongly labeled images such as brain atlases, cropped images, and images thresholded at a very high threshold. We performed meta analyses using the remaining set of curated images with the goal of determining whether results could be obtained using a limited set of unthresholded maps that are similar to results from large coordinate based databases. The analyses focused on two aspects: (i) spatial distribution of activations across all maps (ii) example meta analysis of response inhibition. Code for the analyses is available at https://github.com/NeuroVault/neurovault_analysis.

### Spatial distribution of activations

The goal of this analysis is to explore the spatial distribution of activations across all maps in Neurovault in relation to results previously reported in the literature. The analysis aims to quantify the base rate of activation at each voxel across the entire brain i.e., to identify regions that are activated more or less often across different tasks.

Using coordinate data from the the Neurosynth database, we generated a prior activation probability map based on over 300,000 coordinates drawn from nearly 10,000 published studies. To facilitate fair comparison with the Neurosynth map, we thresholded each map from NeuroVault at a Z or T value of 3 (F maps were excluded). This discretization step approximates the standard Neurosynth procedure of taking discrete peaks reported in studies and convolving them with 3D spheres. We then generated an activation frequency map by counting the proportion of all NeuroVault maps that surpassed the threshold at each voxel.

Figure 4 (middle) shows the NeuroVault frequency map. The distribution is strikingly non uniform throughout gray matter. In particular, the most frequently activated regions include the frontal part of the insula and dorsal anterior cingulate cortex, which form a well known cingulate insulate control network associated with salience processing (Seeley et al., 2007) or maintenance of task sets (ADD: Dosenbach et al, 2006, Neuron). The other structures highlighted in Figure 4 are the inferior parietal sulcus — regions sometimes called the “task positive network” (Fox et al., 2005) — as well as the occipital lobe, encompassing the visual cortex. The presence of the latter likely reflects the fact that the majority of experiments rely on visual stimuli. Interestingly, the networks that are most prominent on this map are largely related to attention and executive control.

**Figure 4.**
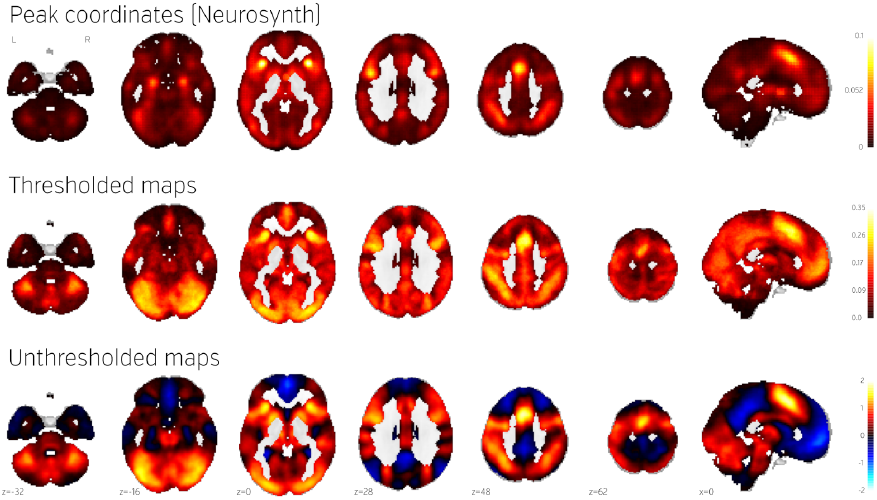
Comparison of frequency of activation across human brain studies obtained using different methods. Top: prior activation probability map obtained from coordinate based meta analysis using NeuroSynth. Middle: proportion of maps in NeuroVault exhibiting values of T or Z higher than 3. Bottom: mean of all T and Z maps (also deposited in NeuroVault). Maps from this figure are available at http://neurovault.org/collections/439/.

The Neurosynth prior activation map is shown for comparison in Figure 4, top. It displays a similar density of activation, with visible attentional networks. It is worth noting that other studies have also reported similar activation density maps (e.g., Nelson et al., 2010). However, the visual cortex is much less present in the Neurosynth map compared to the NeuroVault frequency map. This could potentially be explained by the fact that results NeuroVault includes many statistical maps from fMRI experiments contrasting a single condition with fixation cross baseline. However, most papers report contrasts between conditions removing the effect of the visual stimuli and thus the coordinate database will contain fewer activations in the visual cortex.

We initially thresholded the NeuroVault frequency map in order to facilitate comparison with conventional coordinate based approaches (e.g., the Neurosynth map). However, one important benefit of using unthresholded maps is the retention of additional information in the form of continuous values at all voxels. To investigate what one can gain by using unthresholded maps, we calculated a simple average of all T and Z maps across the entire NeuroVault database (Figure 4, bottom). Unlike the frequency map, as well as the coordinate based meta analysis, this analysis also captures the dominant sign of the activation, accumulating power in regions that may not cross threshold in analyses from individual studies (note that doing a principled statistical inference, e.g. computing a p value or a posterior from this heterogeneous collection of maps would require methodological developments outside of the scope of this article). For example, the average unthresholded map clearly shows regions that respond, on average, by deactivating in the experimental condition relative to the baseline condition (depicted in shades of blue). This pattern spans the default mode network (DMN), which was historically discovered in a similar analysis through observation of consistent decreases in activity across a variety of tasks (Shulman et al., 1997).

### Example Image-Based Meta Analysis using NeuroVault: Response inhibition

To demonstrate how NeuroVault can be used for meta analysis, we turn to the subject of response inhibition. This cognitive concept involves interrupting a prepared or ongoing response to a stimuli as a result of being presented with new information (for review see (Verbruggen and Logan, 2008)). We began by querying the NeuroVault API for statistical maps containing “stop signal” in the task description. This returned 66 maps. We then filtered our set to maps contrasting “stop” and “go” conditions, which resulted in 8 maps across 4 studies (see Table 1). Using the NeuroVault API, we downloaded and visually inspected the maps. Since all of them contained T statistics, we converted them to standardized Z maps prior to the analysis. We estimated the degrees of freedom from the number of participants participating in each study, and this information was also obtained through the NeuroVault API. Since some of the studies contained multiple maps (one study used a test retest protocol, and one used three different variants of the stop signal task) we created one average Z map for each study. We then used Stouffer’s Z score method (Stouffer et al., 1949; Lazar et al., 2002) to combine the results across studies in a fixed effects meta analysis (see Figure 5 top).^3^

**Table 1.**
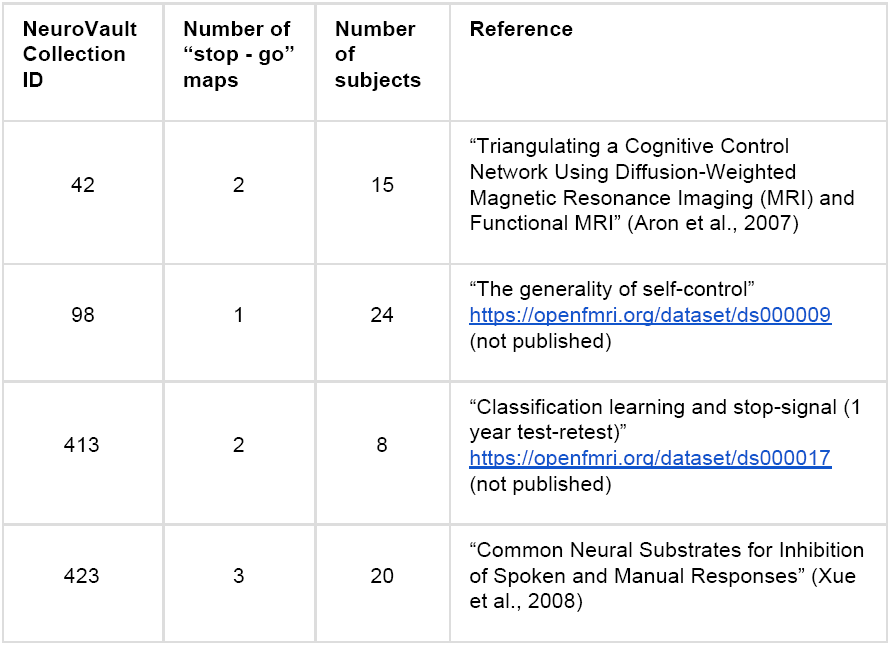
Details of the four studies included in the example meta analysis.

| NeuroVault Collection ID | Number of "stop - go" maps | Number of subjects | Reference |
| --- | --- | --- | --- |
| 42 | 2 | 15 | "Triangulating a Cognitive Control Network Using Diffusion-Weighted Magnetic Resonance Imaging (MRI) and Functional MRI" (Aron et al., 2007) |
| 98 | 1 | 24 | "The generality of self-control"<br>https://openfmri.org/dataset/ds000009<br>(not published) |
| 413 | 2 | 8 | "Classification learning and stop-signal (1 year test-retest)"<br>https://openfmri.org/dataset/ds000017<br>(not published) |
| 423 | 3 | 20 | "Common Neural Substrates for Inhibition of Spoken and Manual Responses" (Xue et al., 2008) |

**Figure 5.**
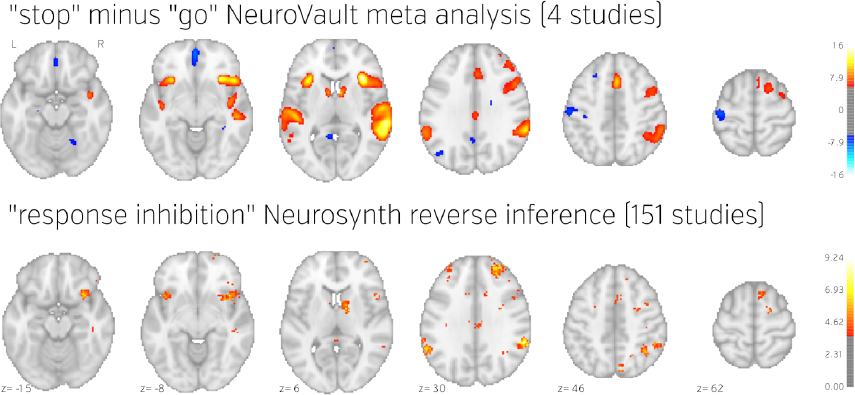
Comparison of image based and coordinate based meta analysis of response inhibition. Meta analysis based on unthresholded statistical maps obtained from NeuroVault (top row) managed to recover the pattern of activation obtained using traditional methods despite including much fewer studies. NeuroVault map has been thresholded at z=6, response inhibition map has been thresholded at z=1.77 (the threshold values were chosen for visualization purposes only, but both are statistically significant at p < 0.05). Unthresholded versions of these maps are available at http://neurovault.org/collections/439/.

The results show consistent activation across the four studies in both left and right inferior frontal gyri and anterior insula as well as left and right parietal cortex. Similar locations have been reported in previous coordinate based meta analyses (Swick et al., 2011; Levy and Wagner, 2011). In contrast to coordinate based meta analyses our analysis also found a deactivation in medial prefrontal cortex. This brain region is one of the hubs of the default mode network, and has been found to be anticorrelated with response inhibition performance (Congdon et al., 2010). This discrepancy is likely caused by the fact that most studies do not report coordinates of deactivation and thus such patterns cannot be picked up by coordinate based meta analyses.

To validate our findings we also compared our results to the “response inhibition” topic map generated by Neurosynth, which is based on 151 studies (see Figure 5). The two maps exhibit remarkable similarity, with the exception of the presence of deactivations and larger cluster extents in the NeuroVault map further validating the notion that an image based meta analysis approach compares favorably to the widely accepted coordinate based approach (cf. Salimi Khorshidi et al, 2009). It is worth noting that our analysis yields plausible results despite being limited to only four studies and a limited number of subjects per study.

## Discussion

We present NeuroVault, a web based platform that allows researchers to store, share, visualize, and decode maps of the human brain. This new resource can improve how human brain mapping experiments are presented, disseminated and reused. Due to its web based implementation NeuroVault does not require any additional software to be installed and thus is very easy to use.

One of the biggest challenges of data sharing platforms is sustainability. Users contributing their data trust that they will be available over an extended period of time. While we cannot make any certain claims about the future, we designed the service in a way to maximize its robustness. NeuroVault is an open source project (the code is available at https://github.com/NeuroVault/NeuroVault) that is dependent only on free and open source components (web servers, content management systems, databases etc.). This means that if the need arises, an individual with minimum web administration experience can set up NeuroVault to run on a new server. Software is not, however, the most important part of the project. To preserve the data we are performing daily offsite backups that are later copied to other locations. The procedure of restoring the service from scratch using the freely available code combined with these backups has been heavily tested. The last component of the service reliability is hardware. It is worth noting that statistical maps take considerably less space than other types of data such as raw fMRI datasets. A 500 GB hard drive (available for $50) can store almost 500,000 statistical maps. Furthermore, the cost of server maintenance and the connection to the Internet can easily be leveraged by existing academic institutions’ infrastructures. In short, we argue that even though no one is able to guarantee long term availability of NeuroVault, due to the nature of its design and the type of data it is dealing with, it is easy and cheap to maintain or host at a new location given there is enough interest and the service will prove to be useful to the scientific community.

NeuroVault is not only a helpful tool for researchers who want to share, visualize and decode their maps, it is also a resource for researchers wanting to perform meta and coactivation analyses. Thanks to the public RESTful API and the CC0 licensing of the data there are no restrictions in terms of how and by whom the data can be used. We hope that this will accelerate progress in the field of human brain imaging and better integrate the growing compendium of resources, as there are many services that could benefit from interaction with NeuroVault. We suggest that Neurosynth and BrainMap can boost the power of their meta analyses by working with unthresholded maps stored in NeuroVault instead of peak coordinates extracted from papers. In our analyses we have showed promising results (replication of Neurosynth frequency map, DMN deactivation and ICA topic maps similar to (Smith et al., 2009)) even with an initial heterogenous set of few maps. The power of an image based meta analysis approach is exemplified by the by the fact that using only a few hundred maps replicated results from much bigger (coordinate based) databases (BrainMap and NeuroSynth cover respectively 2500 and 9000 papers). We are convinced that an increased amount of data will lead to discovering new organizational principles of brain function.

The sharing of neuroimaging data can potentially raise ethical issues related to subject confidentiality (Brakewood & Poldrack, 2013). As NeuroVault is mainly focused on group data analyses, there is little chance that personal information will be included and lead to ethical issues, but the platform allows single subject analysis results to be uploaded. Uploading such data would require researchers to take extra care not to expose the the identity of their subjects.

To minimize the amount of effort needed to create a new collection, the addition of annotated metadata is optional in NeuroVault. Nevertheless, at the users’ discretion, a rich set of metadata can be manually included and stored with the statistical maps. We envision that, in the future, more and more machine readable information will be shared and these metadata will be populated automatically to increase the potential re use of the datasets hosted at NeuroVault. Current efforts (e.g., the previously mentioned BrainSpell), can aid the process of annotating papers (and their corresponding maps) through crowdsourcing. Ideally, machine readable metadata would be made available directly by the software packages used to generate the statistical maps. For example, the NeuroImaging Data Model (NIDM; Keator et al., 2013) is a metadata standard that could be used to withstand metadata loss between an analysis and the upload of the statistical maps into NeuroVault. The NIDM Results standard captures not only the statistic map, but also the design matrix, residuals, group mask and many other pieces of information useful for future analysis. Currently only SPM natively exports to this file format, but we have adopted third party scripts to convert outputs of the FSL analyses (FEAT folders) to NIDM Results on the server side and thus capture richer metadata in a fully automated way, and a solution for AFNI is currently being implemented. To exemplify the importance of such metadata, we present a hypothetical study that aims to train a classifier to predict some outcome from activation maps. It could be the case that effects are due to metadata variables such as the source, software, or scanner, and this finding would only be apparent given that this information is available.

It is also worth pointing out that NeuroVault is not only supporting task based fMRI results. Results from resting state fMRI, PET, VBM, DWI, and most interestingly source reconstructed EEG/MEG experiments can be used with the platform as long as they are NIFTI files in MNI space. We plan to expand this to FreeSurfer surfaces, CIFTI files, and connectomes in the near future. Historically, aggregating results across modalities has been difficult, and we hope that this platform can start to improve upon this situation, by providing one common place for storing and sharing statistical maps.

NeuroVault is also integrated with the Resource Identification Initiative through The Neuroscience Information Framework (NIF, see Gardner et al., 2008 and http://neuinfo.org/). This interdisciplinary project assigns identifiers to resources and tools used in research that are then included in publications and later indexed by Google Scholar and PubMed. These identifiers work with the PubMed LinkOut service (http://www.ncbi.nlm.nih.gov/projects/linkout/) so that links can automatically be made between the tools and publications on web pages describing either. Assigning these resource identifiers to statistical maps, then, would both allow for the creators to track how the maps are used and grant academically acknowledge credit (even in the case when the maps come from unpublished studies).

### Limitations and future directions

One of the biggest limitations of the NeuroVault database is its size and the voluntary nature of data contributions. For any meta analysis to be meaningful the sample of included studies needs to be representative. Including only papers that have corresponding statistical maps in NeuroVault instead of all papers might create unpredictable biases (although this bias is most likely to be towards inclusion of more trustworthy results; see Wicherts et al., 2011). One way of dealing with this is to enforce deposition of statistical maps across all published research. This would be a drastic move, and some data sharing initiatives in neuroimaging in the past were met with considerable opposition from the community (Van Horn and Gazzaniga, 2013). Instead we have reached out to leading journals in the field to encourage (but not require) authors of accepted papers to deposit statistical maps in NeuroVault. So far, NeuroImage, F1000Research and Frontiers in Brain Imaging Methods have joined us in the quest of providing better and more open representation of experimental results. We hope that with time publishing statistical maps will become standard practice.

NeuroVault fills a specific niche in the neuroinformatics ecosystem. The main purpose is to collect, store and share statistical maps. We leave the task of extracting knowledge (tags, labels terms) out of papers and associating them with the statistical maps to other platforms BrainSpell, Neurosynth and BrainMap). We also do not aspire to provide a platform for performing meta analyses (neurosynth and BrainMap facilitate this). This decision is intentional and was made to focus on one specific task and do it well. Thus, in the future we want to focus on i) making the platform more attractive for researchers (so the motivation for data deposition will increase) ii) making the data deposition process easier and automatic extraction of metadata more effective, and iii) reaching out to the community to make sharing of statistical maps a common practice. In terms of the first goal we are working hard on adding new features that will help researchers to understand and visualize their maps. One of such features (currently in beta) is map comparison: users will be able to compare their map with all the other maps deposited in the database and thus easily find experiments with similar imaging results. The second goal will involve tighter integration with the most popular software packages (capitalizing on the NIDM Results standard). We plan to provide a single click solution for uploading maps to NeuroVault that will be available within analysis software such as SPM, FSL and AFNI. Finally the third goal, probably the most important and also the hardest, involves continuous conversations with academic journals and conference organizations such as the OHBM. We hope that by including all of the interested parties in these conversation we will be able to convince the community about the pressing need for sharing statistical maps.

## Conclusion

In this work we have described NeuroVault — a web based repository that allows researchers to store, share, visualize, and decode unthresholded statistical maps of the human brain. This project not only helps individual researchers to disseminate their results and put them in the context of existing literature, but it also enables aggregation of data across studies. Through our analyses we have shown that with only a few hundred statistical maps we can achieve results comparable to those obtained with thousands of sets of coordinates. NeuroVault is free and unencumbered by data use agreements. The data is available and the database queryable via the web interface and RESTful API. This simple and modern platform opens the door to developing novel methods to draw inferences from a meta analytic database.

## Acknowledgements

This work was partially funded by the National Institutes of Health (NIH), R01MH096906 [TY], NSF OCI 1131441 [RP], International Neuroinformatics Coordinating Facility (INCF) and the Max Planck Society [KJG, DSM].

We thank the International Neuroinformatics Coordinating Facility (INCF) Neuroimaging Data Sharing (Nidash) task force members for their input during several discussions.

## Footnotes

1 Collections require only name or DOI fields to be filled. Statistic maps require name, map type (T, Z, F etc.), modality (BOLD fMRI, diffusion, EEG, etc.), and cognitive paradigm (chosen from the list of Tasks in the Cognitive Atlas (Poldrack et al., 2011)) fields to be entered. For more details see Supplementary Tables 1 and 2.

2 BrainSpell is a web platform that allows many users to annotate neuroimaging papers and the results described in them using existing ontologies. It’s based on the crowdsourcing principle anyone is able to contribute their annotations with the assumption that the effort can be spread across multiple people and the consensus will maintain high quality.

3 An alternate approach would be to submit all 8 Z maps to Stouffer’s method, but this neglects the intra study correlation; our practical approach averaging each study’s Zs is conservative but valid. We have presented the results of the analysis of 8 maps here: https://github.com/NeuroVault/neurovault_analysis.

